# A reference model for adult human cardiovascular mechanics

**DOI:** 10.64898/2025.12.10.693559

**Authors:** Timothy W. Secomb, Michael J. Moulton

## Abstract

A previously developed computational model for cardiovascular mechanics is here calibrated using typical values of 22 clinically observed hemodynamic properties in healthy human subjects. The model includes spatially resolved representations of the four heart chambers. The left ventricle is represented as a truncated thick-walled prolate spheroid, with three modes of deformation (short and long axis contraction and torsion). The other chambers are represented as full or partial thick-walled spherical shells. Wave propagation and reflection in the aorta are represented using a one-dimensional model. The closed-loop system is completed using lumped elements representing vascular resistances, compliances and inertances. The resulting system of coupled ordinary differential equations can be solved computationally in less than 0.1 s per cardiac cycle. In the present study, the values of 19 key input parameters to the model are estimated by minimizing the deviation of model predictions from the 22 observed hemodynamic properties. The resulting calibrated reference model provides a baseline for theoretical studies exploring the relationship between fundamental properties of the cardiovascular system, such as ventricular contractility and stiffness or vascular resistance and compliance, and clinically available measurements such as blood pressures, chamber volumes or valve flow waveforms.

## Introduction

The circulatory system consists of multiple components, including the four chambers of the heart and the vessel types comprising the systemic and pulmonary circulations. Normal cardiovascular function is dependent not only on the behavior of the individual components, but also on their interactions. For example, cardiac pumping is strongly dependent on the preload and afterload on the ventricles. The pressure generated in each chamber obviously depends not only on the contractility of the myocardium, but also on the chamber’s size and shape.

As a consequence of these complex interactions among the system components, the underlying causes of circulatory dysfunction can be difficult to determine based on generally available clinical observations. For example, heart failure with preserved ejection function (HFpEF) is considered to be a combination of pathophysiological phenotypes that vary from one patient to another (Borlaug et al., 2023). Contributing factors are believed to include abnormalities in active or passive properties of myocardium, chamber geometry, inflammatory state and fluid loading. However, the presence of these factors and their possible contributions to HFpEF cannot be readily distinguished using non-invasive methods such as echocardiography.

Computational simulations of cardiovascular function provide a potentially useful approach for establishing quantitative relationships between basic properties determining circulatory function and the resulting clinically observable quantities, such as blood pressure, ejection fraction, valve blood flow velocities and waveforms, and chamber dimensions and deformations during the cardiac cycle. Fully three-dimensional models of cardiac function (Davey et al., 2024; Kerckhoffs et al., 2007) are well suited for forward problems, i.e. predicting function based on known cardiac properties and parameters. For the present purpose, however, it is necessary to solve inverse problems, i.e. to deduce the underlying parameters that provide the best fit to observed functional properties. Such parameter estimation requires multiple solutions of the forward problem, and the heavy computational demands of fully three-dimensional models then become a significant limitation. Approaches using simplified cardiac geometries (Lumens et al., 2009; Moulton and Secomb, 2023) are much faster computationally. Estimation of 10 to 20 parameters, requiring 100 to 200 forward solutions, can be achieved within a few minutes.

We (Moulton and Secomb, 2023) have previously presented a closed-loop circulatory system simulation that includes all four chambers of the heart, with systemic and pulmonary circulations. The goal of the present study is to establish a reference set of input parameters for the model, so that it closely represents a set of typical or average values of indices of circulatory function. Such a reference or baseline model is a prerequisite for the eventual use of the model for its intended purpose as a tool for interpretation of non-invasive clinical measurements of circulatory function in humans.

## Methods

### Cardiovascular system properties

For clarity, measurable quantities and indices describing the heart and circulation are here referred to as *properties*, whereas inputs to the theoretical model are referred to as *parameters*. The overall goal of the work is therefore to determine a set of standard properties based on available literature studies, and then to estimate a reference set of parameters such that the predicted properties closely match the literature values.

Cardiovascular properties vary widely in humans with age, sex, state of health and individual environmental and genetic factors. Any single set of properties cannot, of course, describe the variation within a population. The reference model is intended to provide a baseline, which can then be used to examine effects of changes of intrinsic parameters on cardiovascular properties. Here, reference values of cardiovascular properties were derived from several sources. Normal values of pressures in the heart and great vessels were obtained from the Merck Manual (Merck & Co., 2025). End-diastolic and end-systolic chamber volumes were obtained from data for various age groups in adult males and females (Stojanovska et al., 2011; Stojanovska et al., 2014). In that case, the chosen values are averaged over the reported age group and over males and females. Other data sources are described in the following paragraphs. In total, 22 independent properties are available as a basis for parameter estimation.

### Cardiovascular system simulation

The simulation method has been described in detail previously (Moulton and Secomb, 2023). The four-chamber representation of the heart is shown in Figure 1A. The left ventricle (LV) is represented by a thick-walled prolate spheroidal shape, truncated at the base of the heart. In this axisymmetric model, three time-dependent variables describe elongation at constant volume, variation of short-axis length with change of volume, and torsion. The effects of variation of fiber angles through the wall thickness are included. The right ventricle (RV) is represented by a segment of a sphere, and the atria (LA, RA) are represented as spheres. For each of the spherical chambers, a single time-dependent variable describes its deformation. Blood flow through each of the four valves is represented using a model that includes effects of inertia (Mynard et al., 2012). A one-dimensional wave propagation model of the aorta is used, so that effects of wave reflection on the afterload of the LV are included. Other segments of the circulation are represented using lumped elements, including resistances, compliances and inertances (see Figure 1B).

**Figure 1.**
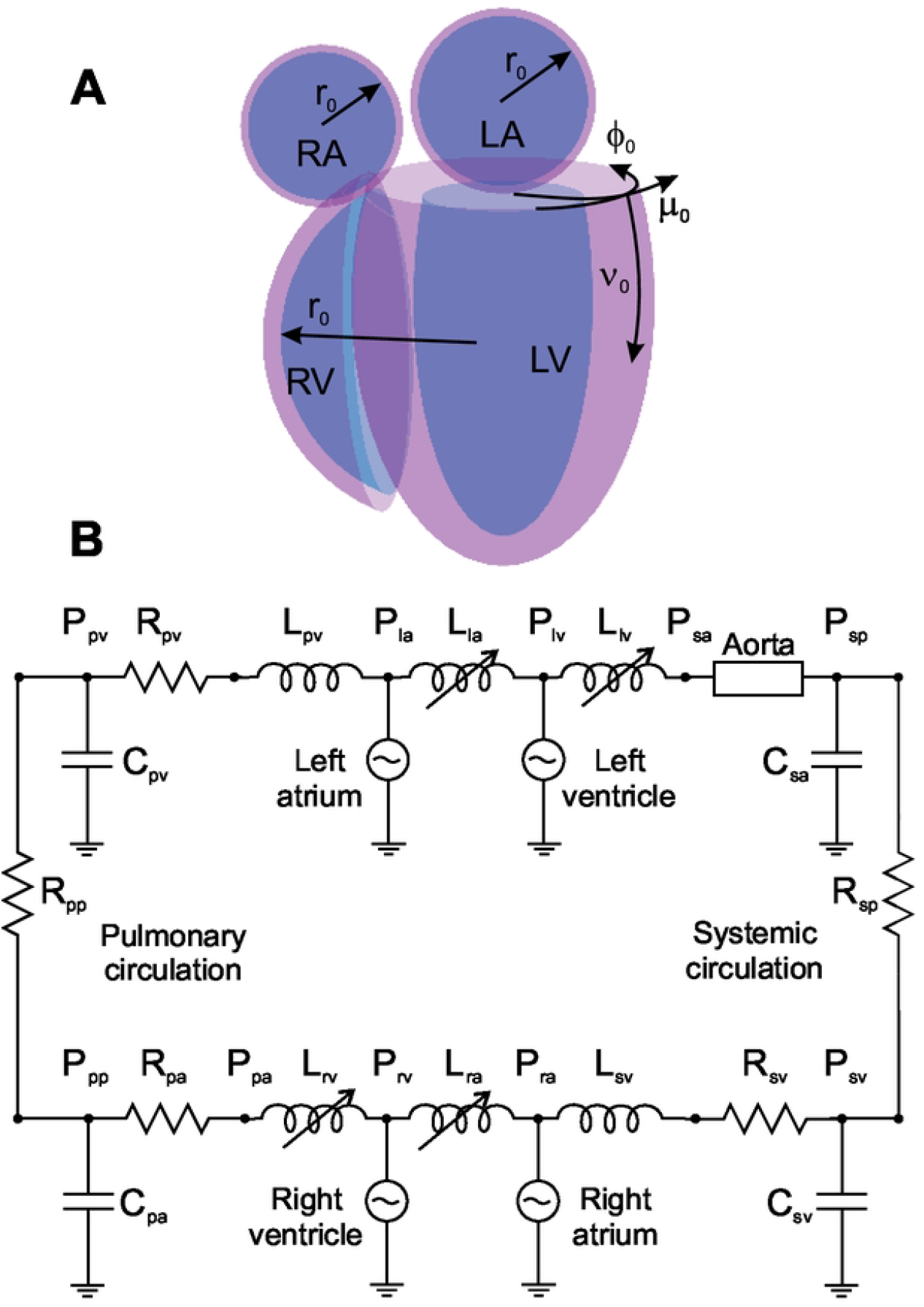
**A.** Schematic view of the four-chamber heart used in the model. Chambers are shown grouped but are mechanically connected only via the heart valves systemic and pulmonary circulations. Reference configurations are shown. The LV is a truncated ellipsoidal shell, described by prolate ellipsoidal coordinates (*μ*_0_, *ν*_0_, *ϕ*_0_). The RV is a part of a spherical shell, and the LA and RA are complete spherical shells, described a radial coordinate *r*_0_. **B**. Schematic of circulatory system model, with components represented by corresponding electrical circuit elements. Inductances: *L*_*la*_, *L*_*lv*_, *L*_*ra*_ and *L*_*rv*_ represent variable inertial effects in valves; *L*_*sv*_ and *L*_*pv*_ represent inertia in major veins. Resistances: *R*_*sp*_ and *R*_*sv*_ represent systemic peripheral and venous vessels; *R*_*pa*_, *R*_*pp*_ and *R*_*pv*_ represent pulmonary arterial, peripheral and venous vessels. Capacitances: *C*_*sa*_ and *C*_*sv*_ represent systemic arterial and venous vessels; *C*_*pa*_ and *C*_*pv*_ represent pulmonary arterial and venous vessels. The aorta is represented by a 1-D wave propagation model.

### Estimation of model parameters

About 80 parameters are needed to completely specify the model (Tables 1 and 2). Other data sources beside the 22 properties mentioned above are therefore needed. Length-tension properties of sarcomeres are based on biophysical observations (ter Keurs et al., 1980). For each chamber, the ratio *k*_*m*_/*k*_*v*_ of the maximum force to the viscous activation-dependent resistance determines the force-velocity characteristics of the muscle. This ratio is fixed at 3 s^−1^ (Gülch and Jacob, 1975). Epi- and endocardial fiber angles in the LV are as observed (Streeter et al., 1969). The valve model was previously calibrated (Mynard et al., 2012). The reference heart rate was obtained from the same source as the chamber volumes (Stojanovska et al., 2014). In that study, healthy subjects were treated with metoprolol if needed, to achieve heart rates of 60 bpm or less, and the resulting average heart rate was 56.9 bpm, implying a cycle time *T*_*c*_ = 1.054 s. Other parameters determining the timing of the cardiac cycle were based on those used previously (Moulton and Secomb, 2023), scaled according to this heart rate.

**Table 1.**
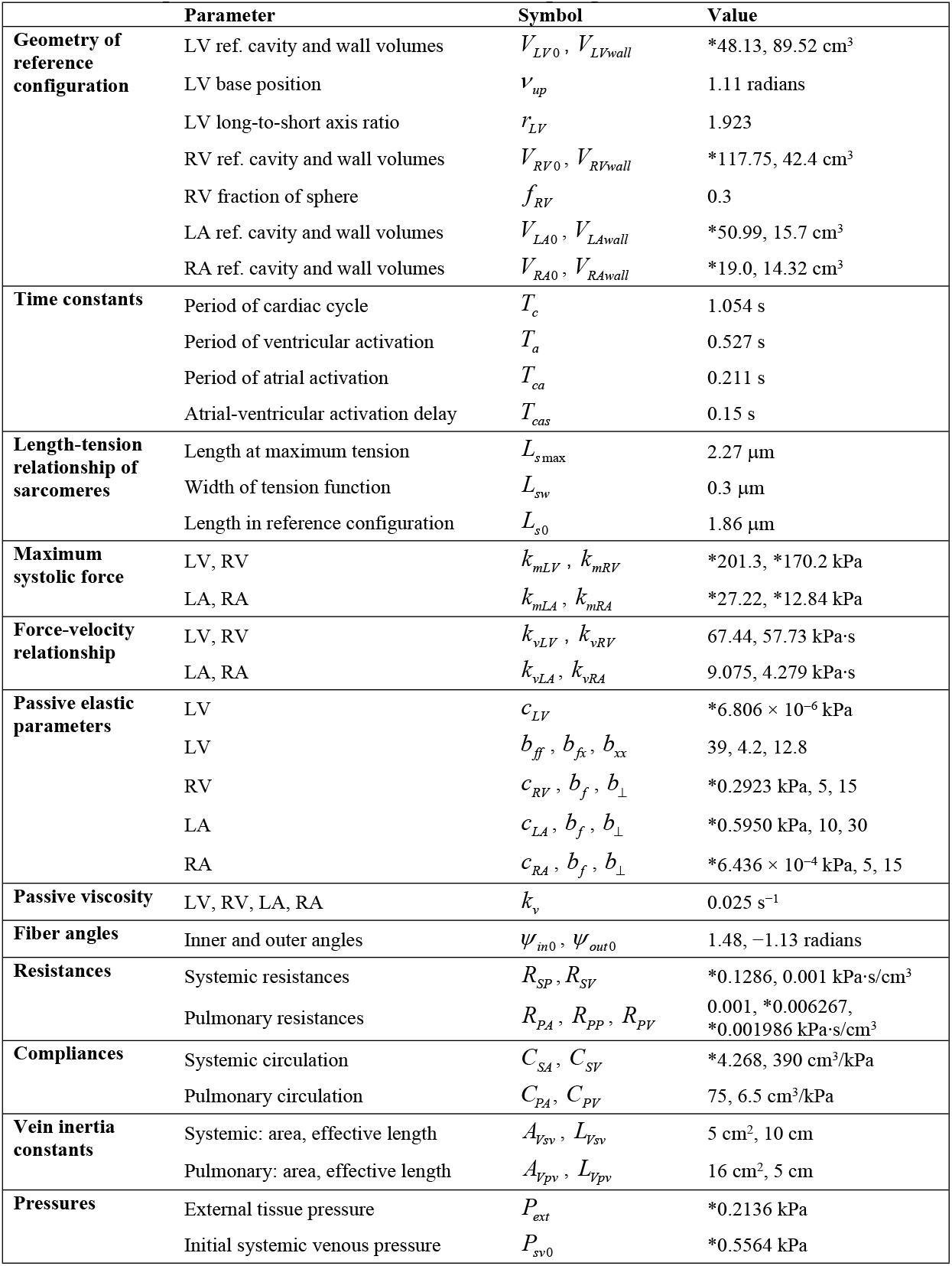
Model parameters: heart chambers and lumped-parameter model. *Fitted value.

**Table 2.** Model parameters: valves and aorta model. *Fitted value.

|  | Parameter | Symbol | Value |
| --- | --- | --- | --- |
| **Valve maximum and minimum orifice areas** | Mitral valve | $A_{mv,open}$ , $A_{mv,closed}$ | 4 cm<sup>2</sup>, 0.001 cm<sup>2</sup> |
| | Tricuspid valve | $A_{tcv,open}$ , $A_{tcv,closed}$ | 4 cm<sup>2</sup>, 0.001 cm<sup>2</sup> |
| | Aortic valve | $A_{aov,open}$ , $A_{aov,closed}$ | 4 cm<sup>2</sup>, 0.001 cm<sup>2</sup> |
| | Pulmonary valve | $A_{pv,open}$ , $A_{pv,closed}$ | 4 cm<sup>2</sup>, 0.001 cm<sup>2</sup> |
| **Valve length constants** | Effective length over which fluid momentum is calculated | $V_{Lmv}$ | 6 cm |
| | | $V_{Ltcv}$ | 5 cm |
| | | $V_{Laov}$ | 1.5 cm |
| | | $V_{Lpv}$ | 1.5 cm |
| **Valve kinetic constants** | Time constants in valve model | $k_{mv,open}$ , $k_{mv,close}$ | 100, 100 (kPa·s)<sup>-1</sup> |
| | | $k_{tcv,open}$ , $k_{tcv,close}$ | 100, 100 (kPa·s)<sup>-1</sup> |
| | | $k_{aov,open}$ , $k_{aov,close}$ | 500, 500 (kPa·s)<sup>-1</sup> |
| | | $k_{pv,open}$ , $k_{pv,close}$ | 500, 1000 (kPa·s)<sup>-1</sup> |
| **Aortic wave model constants** | Compliance of aorta | $G$ | *1.039 × 10<sup>-5</sup> cm<sup>2</sup>/dyn |
| | Taper of aorta | $A_{01}$ | 0.03 cm<sup>-1</sup> |
| | Size of aortic inlet | $A_{00}$ | 4 cm<sup>2</sup> |
| | Length of aorta | $L_{aorta}$ | 50 cm |

Estimates of the volumes of myocardium associated with each chamber wall are used to calculate the camber geometries. For the LV, data (Stojanovska et al., 2014) are averaged over age and sex to yield a reference value *V*_*LVwall*_ = 89.52 cm^3^. For the RV and LA, *V*_*RVwall*_ = 42.4 cm^3^ (Doherty et al., 1992) and *V*_*LAwall*_ = 15.7 cm^3^ (Silva Cunha et al., 2024). For the RA, reported wall volumes of about 30 cm^3^ (Zhao et al., 2023) imply unrealistically large wall thickness. Typical values for RA radius and wall thickness are 1.96 cm (Radiopaedia, 2025) and 0.26 cm (O’Connor et al., 2023), implying *V*_*RAwall*_ = 14.32 cm^3^, a value consistent with other data (Zhan et al., 2024), based on 2 m^2^ body surface area.

LV geometry is described using prolate spheroidal coordinates (*μ, ν, ϕ*), which are related to cartesian coordinates (*x, y, z*) by

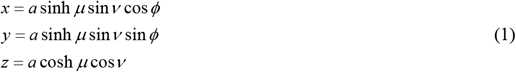

where *a* is a scale factor. The reference configuration is specified by *a* = *a*_0_, *μ* = *μ*_*in*0_ and *μ* = *μ*_*out*0_ on the inner and outer myocardial surfaces, *ν* = *ν*_*up*_ at the base and *ν* = *π* at the apex. The overall shape of the LV is defined in terms of the long- to short-axis ratio of the cavity, *r*_*LV*_. From Eq. (1), *μ*_*in*0_ satisfies

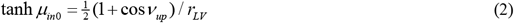

Typical human LV dimensions imply that *ν*_*up*_ = 1.11 and *r*_*LV*_ = 1.923. The volume of a truncated prolate spheroid (0 ≤ *μ* ≤ *μ*_0_, *ν*_*up*_ ≤ *ν* ≤ *π*) can be shown to be

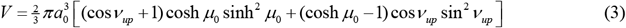

For a given reference chamber volume *V*_*LV*0_, *a*_0_ and *μ*_*out*0_ are calculated by setting *V* = *V*_*LV*0_ and *V* = *V*_*LV*0 +_ *V*_*LVwall*_ respectively. For the other chambers, the reference inner and outer radii satisfy

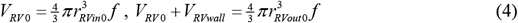

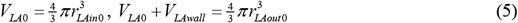

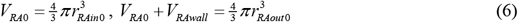

where *f* = 0.3 is the assumed fraction of a spherical shell that represents the RV (see Figure 1A).

The passive elastic properties of the myocardium are described by a transversely isotropic strain- energy function Ψ of exponential type. For the LV,

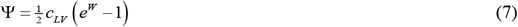

where

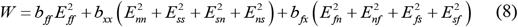

Here, *E*_*ij*_ are the components of Green strain in the fiber coordinates (*s, n, f*) and *c*_*LV*_, *b*_*ff*_, *b*_*xx*_ and *b*_*fx*_ are material parameters (see Supporting Information). For the other chambers, the same form is used, with

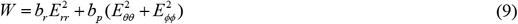

where *E*_*ij*_ are the components of Green strain in spherical coordinates (*r, θ, ϕ*), *b*_*r*_ and *b*_*p*_ are material parameters (see Supporting Information). Published values (Nordbø et al., 2014) are used for the parameters *b*_*ff*_, *b*_*xx*_ and *b*_*fx*_ in the LV. For the other three chambers, lower values for the constants *b*_*r*_ and *b*_*p*_ improve consistency with observations. These nine parameters were not included in the optimization. Another seven parameters (*R*_*SV*_, *R*_*PA*_, *k*_*v*_, *A*_*Vsv*_, *L*_*Vsv*_, *A*_*Vpv*_, *L*_*Vpv*_) are excluded because the properties of interest are weakly dependent on them. Assumed systemic and pulmonary compliances *C*_*SV*_, *C*_*PA*_, *C*_*PV*_ are based on previous estimates (Moulton and Secomb, 2023) and excluded from the optimization. Aortic entrance area, length and taper are independently estimated (Moulton and Secomb, 2023).

### Model formulation and solution

In the model, the dynamic state of the cardiovascular system is described by twenty time-dependent variables. Six variables describe the deformation of the four chambers, six (*a*_1_ to *a*_6_) describe flow rates in and out of the chambers, four describe pressures of the compliant elements in the circulation, and four describe the state of the valves. The conditions for mechanical equilibrium of the chambers are applied by equating the virtual work done on the wall by a small change in each variable *a*_*i*_ to the work done by pressure. Together with the equations describing the lumped-parameter elements and the valve dynamics, these yield a system of twenty ordinary differential equations in time (see Supporting Information), which is solved using a second-order Runge-Kutta scheme, with time steps of about 0.0005 s. Each forward simulation is run for 20 cycles, to allow equilibration of LV and RV stroke volumes. This takes less than 1 s on a laptop computer running at about 3 GHz.

### Parameter estimation

Estimation of the unspecified parameters is performed using a customized version of the Levenberg-Marquardt algorithm. An objective function is defined as

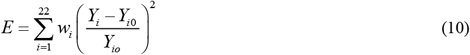

where *Y*_*i*_ is the predicted value of property *i* and *Y*_*i*0_ is its target value. The weighting factors *w*_*i*_ are set to 1, except that increased weights are applied to properties considered to be accurately known and/or of particular importance: LV and RV end-diastolic volumes (*w*_*i*_ = 10), systolic and diastolic blood pressures (*w*_*i*_ = 10), and stroke volume (*w*_*i*_ = 100).

The objective function is minimized with respect to 19 parameters: reference volume (*V*_*LV0*_, *V*_*RV0*_, *V*_*LA0*_, *V*_*RA0*_), maximal active force (*k*_*mLV*_, *k*_*mRV*_, *k*_*mLA*_, *k*_*mRA*_) and passive elastic constant (*c*_*LV*_, *c*_*RV*_, *c*_*LA*_, *c*_*RA*_) for each chamber, flow resistances *R*_*SP*_, *R*_*PP*_, and *R*_*PV*_, compliance *C*_*SA*_, pressures *P*_*ext*_ and *P*_*sv*0_, and aortic compliance *G*. The pressure *P*_*ext*_ is added to all predicted pressures before computing the deviations from targets. This allows for a pressure offset resulting, for instance, from external tissue pressure acting on the heart. The pressure *P*_*sv*0_ is applied as an initial condition to the largest compliance in the system (*C*_*SV*_), thereby controlling total blood volume in the system. Parameter estimation typically requires 100 to 200 forward simulations.

## Results

Tables 1 and 2 list the parameters of the reference model and their values, including those assumed *a priori* and those obtained using the parameter estimation procedure (the latter being indicated by asterisks). Table 3 lists the target values of the model properties based on published data, together with the fitted values obtained by the above-described parameter estimation. The maximum deviation between target and fitted values is less than 10%, and most deviations are less than 5%. Figure 2 illustrates the behavior of key system variables, as predicted by the reference model.

**Table 3.** Target and fitted values of reference hemodynamic properties.

| Property | Unit | Target value | Fitted value | Reference |
| --- | --- | --- | --- | --- |
| LV end-diastolic volume | cm<sup>3</sup> | 151 | 150.90 | (Stojanovska et al., 2014) |
| LV end-systolic volume | cm<sup>3</sup> | 45 | 44.97 | (Stojanovska et al., 2014) |
| LA end-diastolic volume | cm<sup>3</sup> | 80 | 75.55 | (Stojanovska et al., 2011) |
| LA end-systolic volume | cm<sup>3</sup> | 40 | 40.95 | (Stojanovska et al., 2011) |
| RV end-diastolic volume | cm<sup>3</sup> | 181 | 181.34 | (Stojanovska et al., 2014) |
| RV end-systolic volume | cm<sup>3</sup> | 75 | 75.41 | (Stojanovska et al., 2014) |
| RA end-diastolic volume | cm<sup>3</sup> | 52 | 52.28 | (Sun et al., 2023) |
| RA end-systolic volume | cm<sup>3</sup> | 19 | 19.20 | (Sun et al., 2023) |
| Stroke volume | cm<sup>3</sup> | 106 | 105.93 | (Stojanovska et al., 2014) |
| Maximum mitral E-wave velocity | cm/s | 80 | 82.95 | (Eriksen-Volnes et al., 2023) |
| Aorta pulse wave velocity | cm/s | 400 | 384.76 | (Hickson et al., 2010) |
| Mitral E-A ratio |  | 1.4 | 1.30 | (Nagueh et al., 2009) |
| Tricuspid E-A ratio |  | 1.32 | 1.22 | (Shojaeifard et al., 2013) |
| Systolic blood pressure | mmHg | 120 | 119.67 | (Stojanovska et al., 2014) |
| Diastolic blood pressure | mmHg | 73.8 | 73.70 | (Stojanovska et al., 2014) |
| Mean pulmonary vein pressure | mmHg | 9 | 9.06 | (Merck & Co., 2025) |
| Maximum LV pressure | mmHg | 130 | 136.86 | (Merck & Co., 2025) |
| End-diastolic LV pressure | mmHg | 9 | 9.02 | (Merck & Co., 2025) |
| Maximum RV pressure | mmHg | 25 | 26.46 | (Merck & Co., 2025) |
| End-diastolic RV pressure | mmHg | 4 | 4.00 | (Merck & Co., 2025) |
| Mean pulmonary artery pressure | mmHg | 15 | 14.55 | (Merck & Co., 2025) |
| End-diastolic pulmonary artery pressure | mmHg | 9 | 9.00 | (Merck & Co., 2025) |
| Mean right atrial pressure | mmHg | 3 | 3.04 | (Lloyd-Donald et al., 2025) |
| Mean left atrial pressure | mmHg | 8 | 7.56 | (Merck & Co., 2025) |

**Figure 2.**
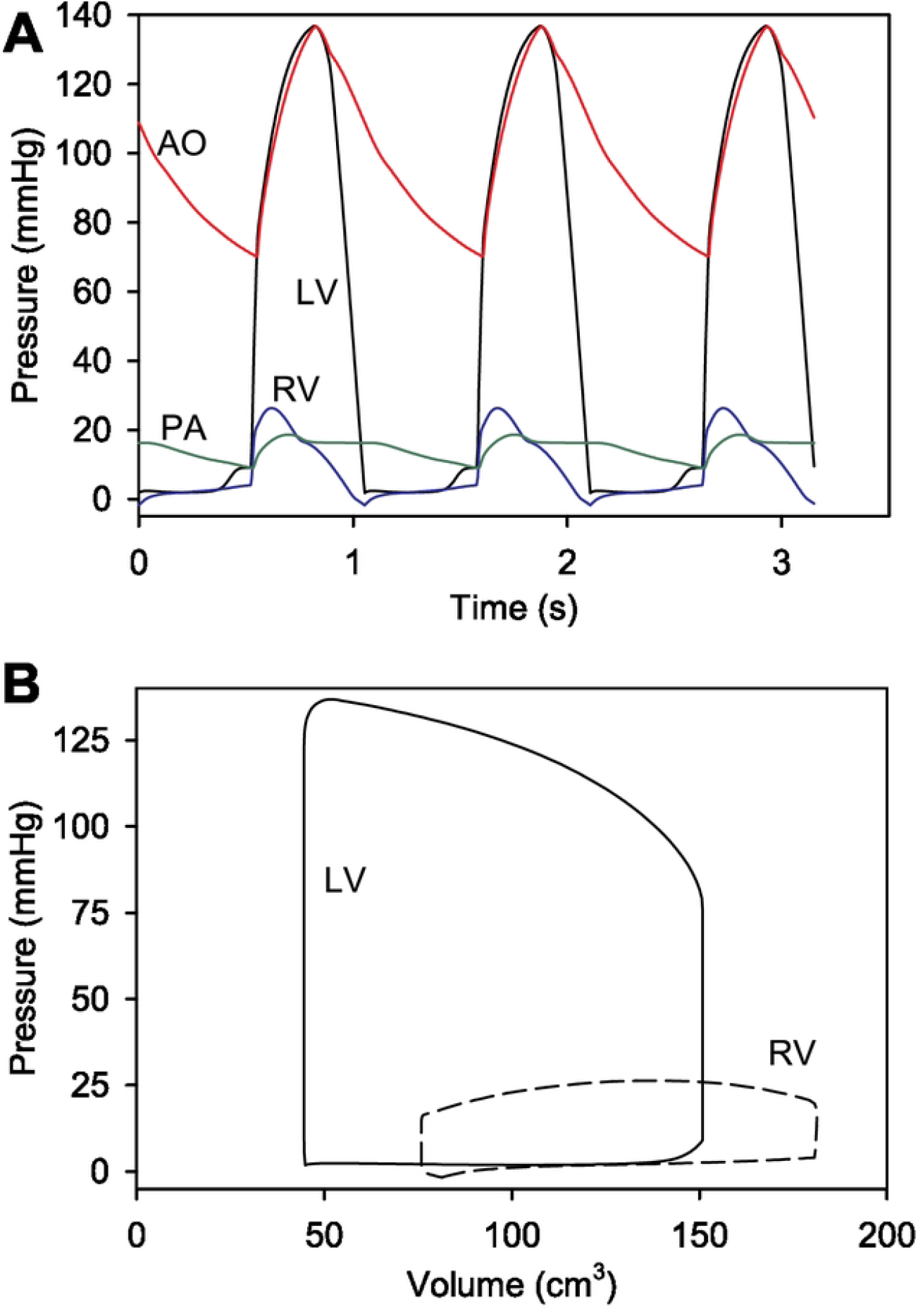
**A.**Predicted variation of pressures during three cardiac cycles. LV: left ventricle. RV: right ventricle. AO: aorta. PA: pulmonary artery. **B**. Pressure-volume loops for the LV and RV.

## Discussion

The objective of the present study is to establish a baseline for theoretical modeling studies on the effects of intrinsic parameters of cardiac and circulatory systems on clinically observable cardiovascular properties. The main result is that our previously developed model (Moulton and Secomb, 2023) is sufficiently detailed and flexible to allow a good match to typical values of 22 clinically observed hemodynamic properties in healthy human subjects. Furthermore, the model allows simulation of 20 cardiac cycles in less than 1 s on a personal computer, allowing solution of inverse problems, i.e. estimation of parameters given observed values of properties, in a few minutes of computer time.

Characteristics of the cardiovascular system vary substantially among individuals, according to sex, age, body mass or surface area, genetic background, state of health, etc. The parameter values estimated here are based on typical observed properties for healthy adults, averaged in some cases over age and sex. Consequently, this reference model does not describe individual characteristics across a human population. However, it could be extended to provide reference models specific for sex, age, and body surface area. Such an extension would utilize measured cardiovascular parameters that are often indexed to body surface area, together with published age- and sex-dependent data (Stojanovska et al., 2011; Stojanovska et al., 2014).

Given the relatively large number of estimated parameters in the model, the optimization process might be expected to yield closer agreement between predicted properties and target values. However, several factors contribute to the lack of precise agreement. The target values are obtained from several different sources and are subject to measurement errors, so they may not be internally consistent. Also, the *a priori* parameter values may be inaccurate, and the model is of course a simplified representation of the actual system.

A further challenge in parameter estimation is that widely covarying values of two or more parameters may yield only slight changes in predicted model properties. In the present model, this is notably the case for the reference cavity volume (*V*_*LV*0_, for example) and the multiplicative constant in the strain-energy function (*c*_*LV*_, for example). Because strains are computed relative to the reference cavity shape, a change in the reference volume leads to a shift in the Green strains *E*_*ij*_. A multiplicative change in the constant *c*_*LV*_ can approximately compensate for this shift, leading to almost the same diastolic filling curve. The large values of the constants *b*_*ff*_, etc., in equations (8) and (9) result in wide variations among the estimated values of *c*_*LV*_, *c*_*RV*_, *c*_*LA*_ and *c*_*RA*_ (see Table 1). These values do not imply wide variations in the actual passive elastic properties of the chambers, but are a consequence of the lack of independent data on the reference or unloaded volumes of the chambers.

In summary, a computationally efficient, mechanistic model of the human circulatory system, including spatially resolved heart chambers, is here calibrated using typical or average values of frequently measured system properties. The resulting reference model can be used as a basis for a range of further investigations, including predicting effects of altered system parameters, estimation of underlying parameters based on clinically observable cardiovascular properties, and development of patient-specific models.

